# Microtubules and actin filaments direct nuclear movement during the polarisation of *Marchantia* spore cells

**DOI:** 10.1101/2024.02.23.581750

**Authors:** Sarah Attrill, Liam Dolan

## Abstract

Multicellular organisms typically develop from single cells, the polarity of which establishes the first body axis of the organism. The multicellular haploid stage of land plants develops from a single haploid cell produced by meiosis – the spore. Starting from a non-polar state, these spores develop polarity and divide asymmetrically to establish the first apical-basal axis of the plant body. In the spore of the liverwort, *Marchantia polymorpha*, we show that the nucleus migrates from the cell centroid to the side of the cell to define the future basal pole. A microtubule organising centre leads this migration by initiating a dense microtubules array towards the cortex at the basal pole. Simultaneously, cortical microtubules disappear from the apical hemisphere but persist near the basal pole. A dense network of fine actin filaments also accumulates between the nucleus and the basal cell cortex. These data demonstrate that microtubules and actin filaments reorganise during the polarisation of the *M. polymorpha* spore. We speculate that signals orient microtubules and actin filaments during spore polarisation, resulting in the formation of a fine actin filament network between the nucleus and cell cortex that moves the nucleus to the future basal pole.

**SUMMARY STATEMENT:** Microtubules and actin filament dynamics are required for the basal movement of the nucleus which establishes cell asymmetry before cell division in the *Marchantia* spore.

## INTRODUCTION

Polarisation – defined as the development of asymmetry across a cell – is an essential process in the development of multicellular organisms. There are two multicellular stages in the life cycle of land plants. The multicellular diploid phase is derived from the zygote produced by fertilization of the egg cell. The multicellular haploid phase is derived from the spore cell produced by meiosis. The polarity of each cell type – zygote and spore – defines the orientation of the plants’ apical-basal body axis. Zygote polarity is inherited from the egg cell (Faure et al., 2002). By contrast, there is no evidence of inherent polarity in spores (Roeder et al., 2022). Rather, spores start in a non-polar state and polarity develops *de novo* within days of germination. Here we test if microtubules and actin filaments are involved in the establishment of asymmetry in spores.

Microtubules and actin filaments reorganise during the polarisation of animal zygotes. In *Drosophila melanogaster*, the orientation of microtubules directs the movement of the nucleus to one cell pole, polarising distinct activities to opposite ends of the cell (Bernard et al., 2018; Huynh and St Johnston, 2004). While in *Caenorhabditis elegans,* the site of sperm entry sets in motion events that reorganise the actin cortex (Munro and Bowerman, 2009). Subsequently, on actin-myosin contraction, cortical flows are generated which direct polarity determinants to the cell poles (Munro et al., 2004). In both examples, this polarity defines the future anterior-posterior axis of the animal. In the zygote of *Arabidopsis thaliana*, the future apical-basal body axis is defined by microtubules and actin filaments which direct cell elongation and nuclear migration respectively (Kimata et al., 2016; Kimata et al., 2019). Given that microtubules and actin filaments play key roles in the polarisation of fly, worm, and angiosperm zygotes, we hypothesised that the cytoskeleton would also function in spore polarisation.

The multicellular haploid phase (gametophyte) of non-seed land plants – bryophytes, lycophytes and monilophytes – begin life as a spore; a single haploid cell produced by meiosis and encased in a sporopollenin wall. Bryophyte spores expand upon germination and divide to form a multicellular structure known as a prothallus (if growth is in two or three dimensions) or a protonema (if a one-dimensional filament forms). The spore of *Marchantia polymorpha* divides asymmetrically to form a large apical cell which proliferates, and a small basal cell which differentiates into a rhizoid (O’Hanlon, 1926; Shimamura, 2016). Since the first division is asymmetric, this indicates that apical-basal polarity is established in the spore before cell division. Very little is known of the mechanisms driving this polarity, except that two microtubule organising centres – known as polar organisers - and their associated arrays migrate to one cell pole prior to cell division (Sakai et al., 2022). We hypothesised that both microtubules and actin filaments are required to polarise the spore of *M. polymorpha*.

Here, we show that the nucleus moves from the centroid to one side of the spore cell, establishing the future basal pole and setting up the asymmetry of the first division. Nuclear migration is led by a polar organiser nucleating a dense astral array that polymerises towards the cortex at the basal pole. Concurrently with nuclear movement, cortical microtubules disappear from the apical hemisphere but remain near the basal pole. Furthermore, a network of fine actin filaments forms between the nucleus and the basal cell cortex. The data indicate that microtubules and actin filaments together coordinate nuclear migration and the establishment of polarity in the germinating spore of *M. polymorpha*.

## RESULTS

### *Marchantia polymorpha* spores divide asymmetrically between 24 and 32 hours after plating on nutrient media

The multicellular, haploid phase of the *Marchantia polymorpha* life cycle is derived from a spore. To define the stages of spore development, germinating spores resulting from a cross between wildtype Tak-1 and Tak-2 were imaged at 24, 32, 48 and 72 h after plating (Fig. 1A, Fig. S1A, B). At 24 h, all spores were single-celled and had chloroplasts. By 32 h, most cells in the population had divided asymmetrically forming a large chloroplast-filled apical cell and a small basal cell with relatively fewer chloroplasts. By 48 h, the basal cell had elongated to form a short rhizoid and the apical cell had divided. By 72 h, the apical cell had undergone multiple rounds of cell division, while the basal rhizoid cell had elongated further but not divided. In conclusion, spore populations develop highly asynchronously with most spores dividing between 24 h and 32 h after plating.

**Fig. 1:**
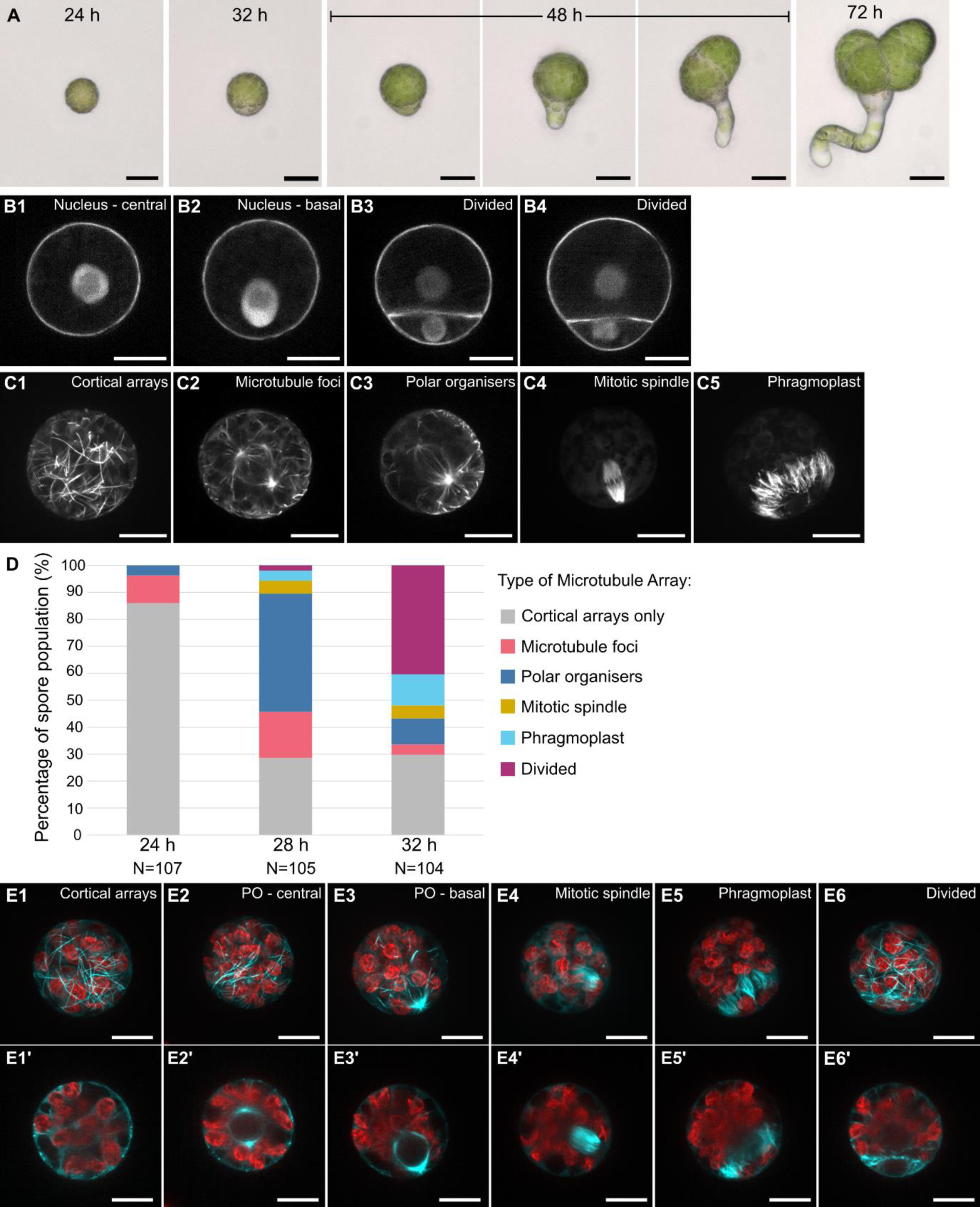
*Marchantia polymorpha* spores divide asymmetrically between 24 and 32 hours after plating and microtubule organisation progressively changes. (A) Development of wild type spores at 24, 32, 48 and 72 h after plating. Each spore is a different individual from the same population. Scale bars, 20 μm. (B) Position of the nucleus and division plane in spores expressing *p*Mp*ROP:mScarletI-N7 - p*Mp*UBE2:mScarletI-*At*LTI6b* at 30 h after plating. Presented are central slices in XY of four spores. Scale bars, 10 μm. (C) Microtubule organisation in spores expressing *p*Mp*EF1α:GFP-*Mp*TUB1* at 29 h after plating. Presented are full Z-projections (1, 4, 5) and Z-projections of central slices (2, 3) of five spores. Scale bars, 10 μm. (D) Percentage of the spore population with each type of microtubule array at 24, 28 and 32 h after plating. N is the number of spores analysed within the population at each timepoint. (E) Chloroplast (red) and microtubule (cyan) organisation in spores expressing *p*Mp*EF1α:GFP-*Mp*TUB1* at 29 h after plating. Presented are Z-projections and central slices in XY (‘) of six spores. PO refers to polar organisers.

Since the first division of the spore is asymmetric, we hypothesised that the nucleus would be positioned asymmetrically within the cell before mitosis. To visualise the position of the nucleus in spores, we imaged spores expressing a nuclear reporter *p*Mp*ROP:mScarletI-N7* (mScarletI-N7) and a plasma membrane reporter *p*Mp*UBE2:mScarletI-*At*LTI6b* (mScarletI-AtLTI6b) (Mulvey and Dolan, 2023; Sauret-Güeto et al., 2020). In single-celled spores at 29 h after plating, the nucleus was either located near the geometric centre of the spore (the centroid) or near one pole of the cell (Fig. 1B1, 2). In two-celled spores the division plane was positioned highly asymmetrically, and each daughter cell had a central nucleus (Fig. 1B3, 4). These data confirm the asymmetry of the first division and suggest that this asymmetry results from the nucleus moving from the cell centroid to one pole of the cell.

### Microtubule organisation changes during the development of spore asymmetry

We hypothesised that a cytoskeleton-based mechanism would be involved in directing the nucleus from the cell centroid to one pole of the cell. We characterised the organisation of microtubules during early spore development by labelling live microtubules with the *p*Mp*EF1α:GFP-*Mp*TUB1* (GFP-MpTUB1) reporter (Buschmann *et al*., 2016). Five distinct microtubule arrays have previously been described in the epidermal cells of *M. polymorpha* thalli; an interphase cortical array located next to the cell membrane (Fig. S1C1); microtubule foci that form on the nucleus surface in preprophase (Fig. S1C2); perinuclear and astral arrays nucleated from two microtubule organising centres - termed ‘polar organisers’ - located on opposite sides of the preprophase nucleus (Fig. S1C3); mitotic spindle arrays (Fig. S1C4); and phragmoplast arrays for cytokinesis (Fig. S1C5). These five microtubule arrays were also observed among populations of unsynchronised spores at 29 h after plating (Fig. 1C). Microtubule foci were often centrally positioned in spores. Whereas the mitotic spindle and phragmoplast were consistently positioned near one pole of the cell. Chloroplasts were densely populated in the hemisphere opposite to where the mitotic spindle were positioned (Fig. 1E). The position of polar organisers varied between the cell centroid and one pole. In summary, the same microtubule arrays are present in spores and epidermal cells, but mitotic and cytokinetic arrays are polar localised in spores.

To determine the timing of each microtubule reorganisation event, spores were imaged at 24, 28 and 32 h after plating and their microtubule organisations quantified (Fig. 1D, Fig. S1D). Cortical arrays were present in all spores at 24 h. Microtubule foci and polar organisers first appeared in the population at 24 h, and by 28 h they were present in 61% of spores. The first cell divisions occurred at 28 h, and by 32 h almost 60% of the population were either in mitosis, cytokinesis or had divided. These data indicate that the microtubule organisation progressively changes – from cortical microtubule arrays to preprophase arrays to mitotic spindle and phragmoplast arrays – in the spore population between 24 h and 32 h after plating as the population develops asynchronously.

### Polar organisers and the nucleus move from the cell centroid to the basal pole before cell division

To determine the exact order of microtubule reorganisation events in spores, we analysed the microtubule dynamics in individual spores by timelapse microscopy. First, we investigated polar organiser formation. Microtubule foci and polar organisers were present in a small percentage of spores at 24 h. By 28 h, almost half the spore population formed polar organisers (Fig. 1D). Polar organisers are therefore not inherited from the sporocyte (spore mother cell) but are formed *de novo* after spore germination. Timelapse imaging of single spores at 29 h after plating demonstrated that multiple low intensity foci coalesce to form the two brighter foci – polar organisers – on opposite sides of the nucleus (Fig. 2A). This is consistent with two polar organisers forming through aggregation of multiple smaller foci, as previously observed in *M. polymorpha* tissue cells (Buschmann *et al.,* 2016).

**Fig. 2:**
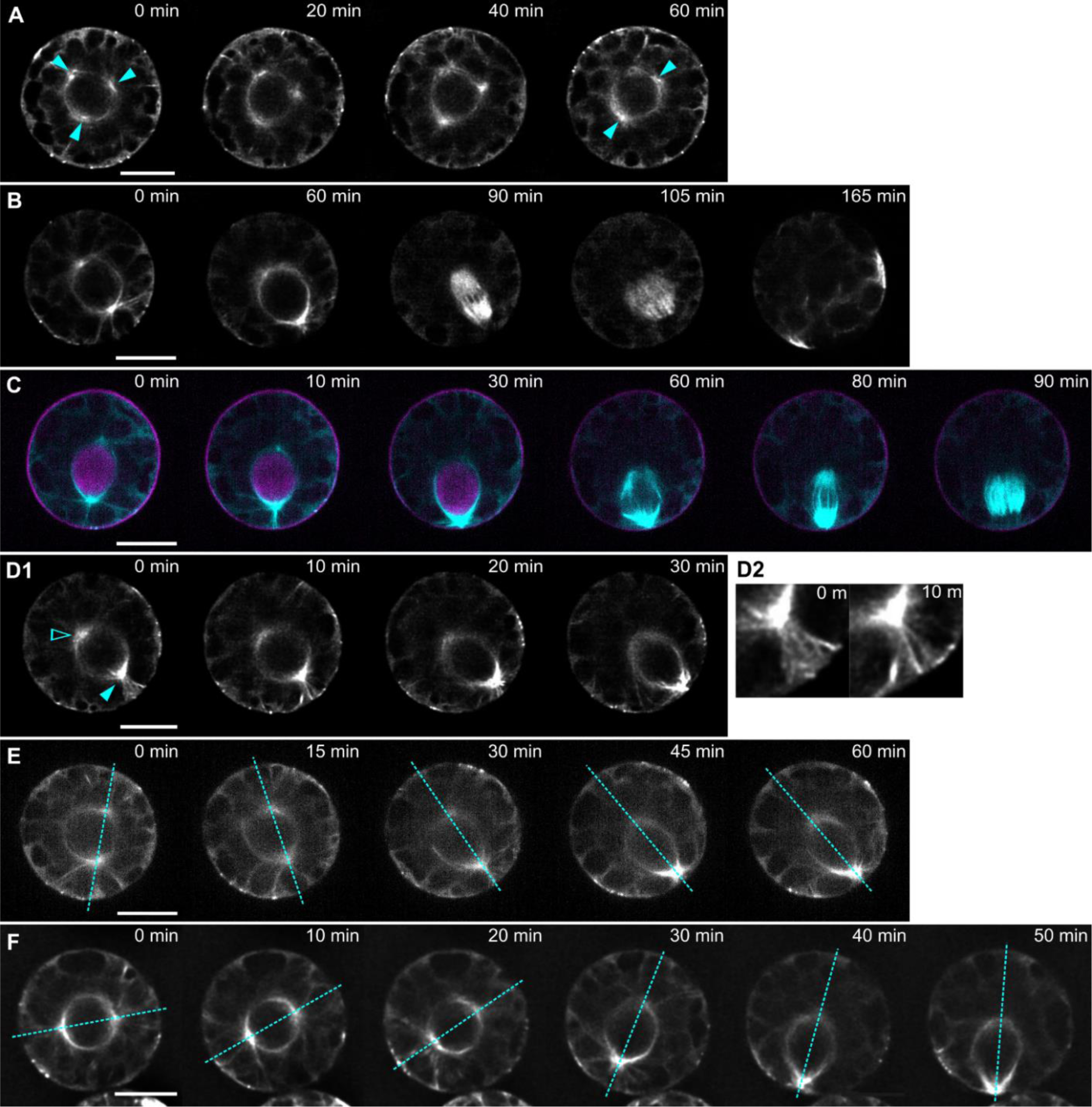
Migration of the nucleus from the cell centroid to the basal pole is led by a polar organiser and dense astral array. Timelapses of microtubule reorganisation in spores expressing *p*Mp*EF1α:GFP-*Mp*TUB1* starting at 29 h after plating. Presented are central slices in XY. Images (A, B, D, F) were deconvolved. Scale bars, 10 μm. (A) Formation of two polar organisers (arrows at 60 min) from three foci (arrows at 0 min). (B) Major microtubule reorganisation events during spore development: polar organiser migration (0 to 60 min), mitosis (90 min) and cytokinesis (105 to 165 min). (C) Co-migration of the nucleus (magenta) and polar organisers/microtubules (cyan) from the cell centroid to the basal pole followed by mitosis. Nuclear position was captured by the expression of *p*Mp*ROP:mScarletI-N7 - p*Mp*UBE2:mScarletI-*At*LTI6b*. (D) A dense astral array spans between the basal polar organiser (filled arrow) and basal cortex. This array is magnified in D2. There is no such array between the apical polar organiser (empty arrow) and apical cortex. (E) Subtle rotation of the polar organiser axis (dotted line) before/during migration to the basal pole. (F) Large 90° rotation of the polar organiser axis (dotted line) before/during migration to the basal pole.

To investigate the events after polar organiser formation, further timelapses of individual spores were captured at 29 h after plating. Polar organisers and perinuclear arrays were initially located near the cell centroid, but 60 min later were positioned near one cell pole (Fig. 2B). By 90 min this structure had disassembled and was replaced by the mitotic spindle. By 105 min the mitotic spindle was replaced by a phragmoplast aligned with the spindle equator. Cytokinesis was completed at this plane between 105 and 165 min. In summary, the polar organisers and associated perinuclear arrays moved from the cell centroid to one cell pole, where the future mitotic spindle and phragmoplast formed. This aligns with previous observations made by Sakai et al., 2022. We define this pole as the ‘basal pole’ – as this domain will form the basal cell on division – and the opposite pole as the ‘apical pole’.

Since the nucleus and polar organisers both localise to the basal pole before cell division, we predicted that these structures would move together. To define the position of the nucleus and polar organisers simultaneously, spores expressing the double nuclear-plasma membrane reporter, mScarletI-N7-mScarletI-AtLTI6b, and the microtubule reporter, GFP-MpTUB1, were imaged at 29 h after plating. At the first timepoint, the nucleus localised between the two polar organisers in the cell centroid (Fig. 2C). As the polar organisers migrated from the cell centroid to the basal pole, the nucleus also migrated. The nuclear signal disappeared at 60 min, indicating disintegration of nuclear envelope, followed by appearance of the mitotic spindle. During cytokinesis, two daughter nuclei were reformed (Fig. S3A, B). In summary, the nucleus and polar organisers migrate together to the basal pole before the first cell division.

Next, we investigated the dynamics of the two polar organisers and the associated astral arrays during nuclear migration. In timelapses, one polar organiser was consistently positioned closer to the cell cortex and appeared to lead the migration (Fig. 2D1). A dense array of astral microtubules always extended from this leading polar organiser to the cell cortex (Fig. 2D2). This cortical region at the basal pole is defined as the ‘basal cortex’ and the leading polar organiser as the ‘basal polar organiser’. During migration, the opposing ‘apical polar organiser’ initiated few astral microtubules which extended out to the ‘apical cortex’. These data indicate that the formation of a dense astral array - initiated from the basal polar organiser and polymerising towards the basal cortex - is correlated with nuclear migration.

Pairs of polar organisers on opposite sides of the nucleus often rotated before and during their migration to the basal cortex. In timelapses, the polar organiser axis – the straight line joining the two polar organisers – rotated around the cell centroid (0 to 30 min in Fig. 2E, F). The degree of rotation varied between spores; often the shift was subtle (Fig. 2E) but occasionally larger rotations of up to 90° from the starting orientation were observed (Fig. 2F, Fig. S3C). This initial rotation roughly aligned the polar organiser axis parallel to the future mitotic spindle axis. On migration the polar organiser axis continued to rotate, by a smaller degree, to fully align with the future apical basal axis. This rotation suggests that polar organisers orient and migrate towards a specific, pre-determined cortical region.

### An asymmetric distribution of cortical microtubules develops progressively as the polar organisers migrate towards the basal pole

Cortical microtubules form a network throughout the cell cortex that reorganises during spore polarisation. At 24 h after plating, before polar organiser formation, a sparse random network of microtubules covered the entire spore cortex (Fig. 3A). Most cortical microtubules were short and highly dynamic, treadmilling across the spore cortex (Fig. S3D). This random organisation was also present in spores at 29 h after plating, when microtubule foci or polar organisers were positioned near the cell centroid (Fig. 3B, C). By contrast, when the basal polar organiser was positioned at the basal cortex, cortical microtubules were consistently dense in the basal hemisphere but absent from the apical hemisphere (Fig. 3D). In summary, cortical microtubules are randomly organised before and during polar organiser formation. Subsequently, after polar organiser migration, they disappear from the apical hemisphere and accumulate near the basal pole.

**Fig. 3:**
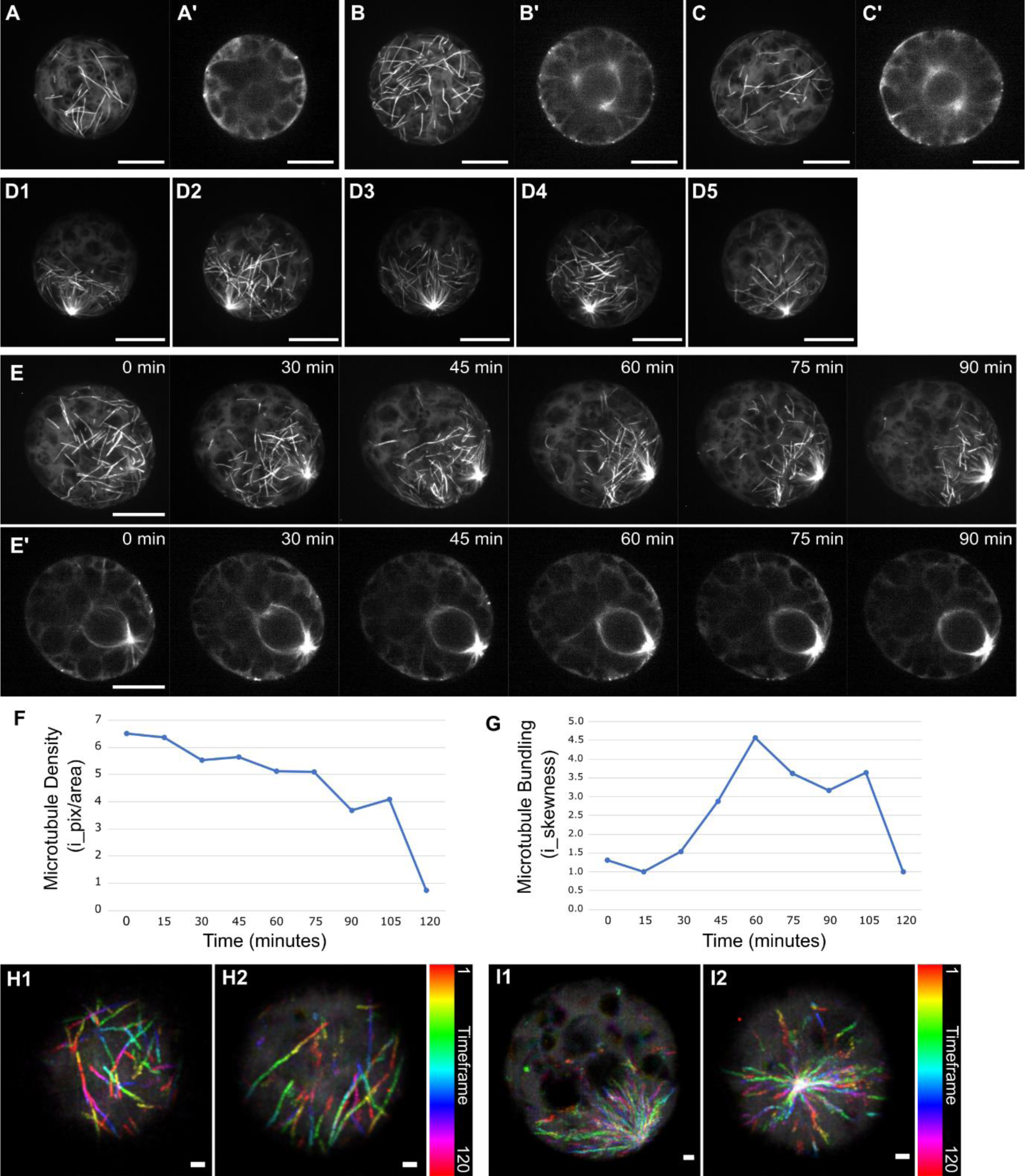
Cortical microtubules deplete from the apical hemisphere during polar organiser migration to the basal pole. (A - D) Organisation of cortical microtubules in spores expressing *p*Mp*EF1α:GFP-*Mp*TUB1* at 24 h (A) or 29 h (B, C, D) after plating. Presented are Z-projections and central planes in XY (‘). Cortical arrays were randomly arranged in spores with no cytosolic arrays (A), microtubule foci (B) and polar organisers near the cell centroid (C). Cortical arrays were basally localised in spores with polar organisers at the basal pole (D1-5). Scale bars, 10 μm. (E) Timelapse of a spore expressing *p*Mp*EF1α:GFP-*Mp*TUB1* starting at 29 h after plating. Presented are Z-projections showing the depletion of cortical microtubules from the apical cortex as the polar organiser, seen in the central XY planes (’), migrates to the basal pole. Scale bars, 10 μm. (F) Plot of the cortical microtubule density in spore E over 120 min. (G) Plot of the cortical microtubule bundling in spore E over 120 min. (H - I) Tracking of microtubule plus ends in spores expressing *p*Mp*EF1α:GFP-*At*EB1a* at 29 h after plating. Presented are colour-coded temporal projections of GFP-EB1 in the cortex (H) and near the basal polar organiser (I) in two spores each. Timelapses were captured at 1.2 s intervals. Images were deconvolved and Z-projected before temporal projection. Scale bars, 1 μm.

We correlated cortical microtubule reorganisation to polar organiser migration through timelapse microscopy of spores at 29 h after plating. As the polar organiser moved to, and remained at, the basal pole, the cortical microtubule network progressively decreased in density (20 to 50 min in Fig. 3E, 60 to 90 min in Fig. S3E). This depletion started at the apical pole and proceeded towards the basal pole. Eventually only a few short, scattered microtubules remained in the basal hemisphere (60 min in Fig. 3E). Quantification showed that cortical microtubule density progressively decreased overtime (Fig. 3F). By contrast, microtubules became increasingly bundled overtime, peaking once the polar organiser had docked at the basal pole (60 min in Fig. 3G). Both measures then dropped when the remaining cortical microtubules disappeared at mitosis (120 min in Fig. 3F, G). In conclusion, there is a gradual change from a random to an asymmetric distribution of cortical arrays as polar organisers migrate to the basal pole.

We hypothesised that the asymmetry of the cortical arrays could be linked to microtubule depolymerisation in the apical hemisphere and/or increased initiation of microtubules from the basal polar organiser which feed into the basal hemisphere. To determine if cortical microtubules can be initiated from polar organisers, polymerising microtubules were tracked using the END BINDING 1 (EB1) protein which binds to microtubule plus ends (Chan et al., 2003). Timelapses of spores expressing the *p*Mp*EF1α:GFP-*At*EB1a* reporter (GFP-AtEB1) were captured at 1.2 s intervals starting at 29 h after plating (Buschmann *et al*., 2016). Z-projections of each timepoint were coloured-coded and overlaid to generate temporal projections. At the cortex, EB1 comets moved in distinct tracks confirming that GFP-AtEB1 labelled polymerising cortical microtubule ends in spores (Fig. 3H; Movie 1). In some spores, a high density of EB1 comets rapidly moved away from a single point (a polar organiser) (Fig. 3I; Movie 2). This reflects a high polymerisation rate of microtubules initiating from the basal polar organiser. Tracking of EB1 comets indicated that some microtubules continued to polymerise into the cell cortex. This is consistent with the hypothesis that the cortical microtubule population in the basal hemisphere is, at least in part, maintained by microtubules initiated from the basal polar organiser.

### The final position of the basal polar organiser and nucleus defines division asymmetry

We hypothesised that polarity would be fixed once the polar organisers had migrated to the basal pole. The polar organiser axis frequently rotated before migration (Fig. 2E, F). However, after migration and docking of the basal organiser at the basal pole, no further rotation was ever observed. Instead, the polar organisers remained in this docked position for a considerable time before their disintegration, as indicated by the change in fluorescence signal from distinct points to broad flat areas around the perinuclear arrays (30 to 40 min in Fig. 4A). In summary, docked polar organisers do not move or reorient. This is consistent with the hypothesis that apical-basal polarity is fixed once the nucleus is at the basal pole.

**Fig. 4:**
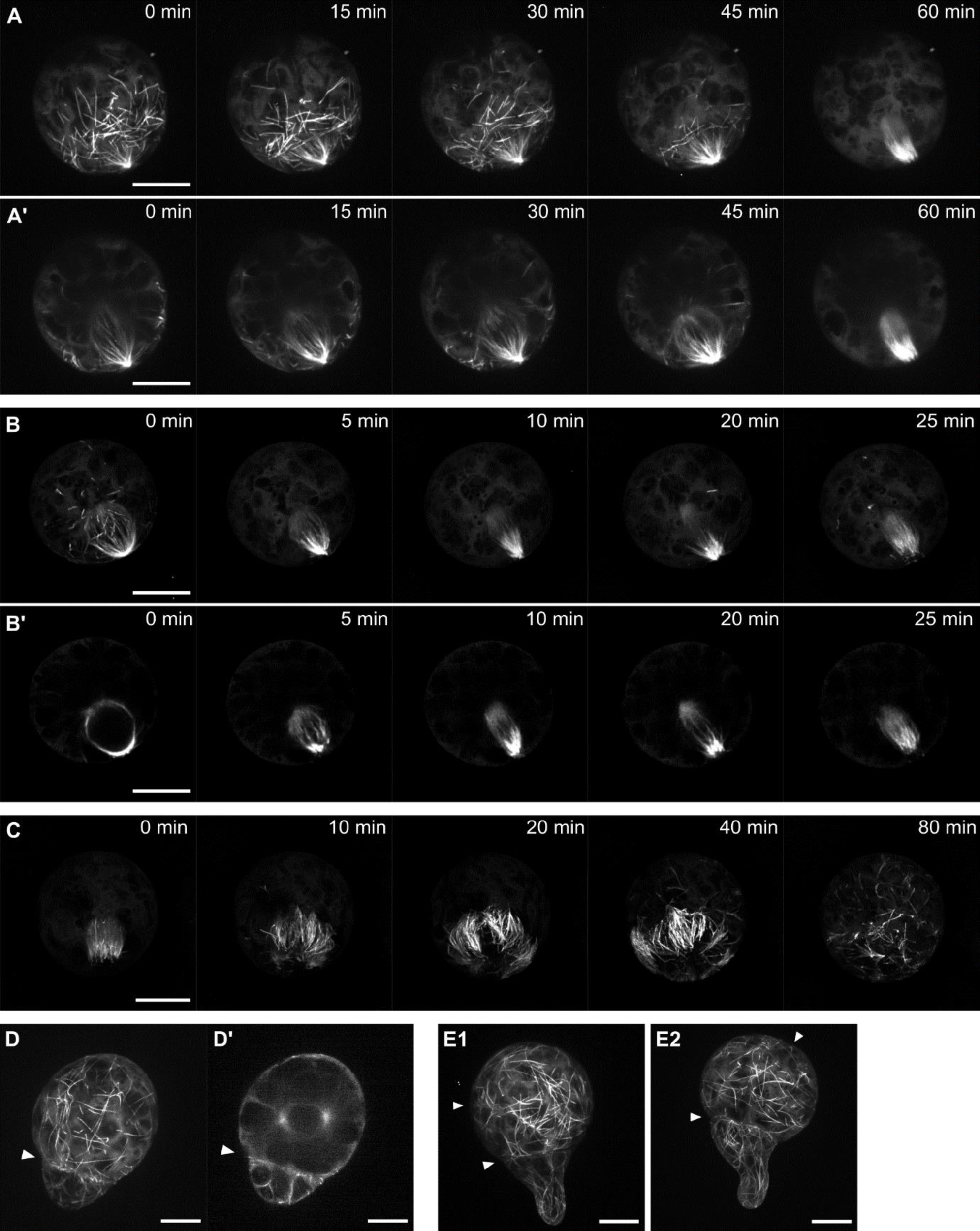
The position of the basal polar organiser defines the first asymmetric division plane. (A - C) Timelapses of microtubule reorganisation before and during cell division in spores expressing *p*Mp*EF1α:GFP-*Mp*TUB1* starting at 29 h after plating. Timelapses show the persistence of the polar organiser at the basal pole and its breakdown (A), mitosis (B) and phragmoplast expansion (C). Presented are Z-projections and central planes in XY (’). Images B, C were deconvolved. Scale bars, 10 μm. (D, E) Microtubule organisation in multi-celled sporelings expressing *p*Mp*EF1α:GFP-* Mp*TUB1* at 50 h after plating. Images show a 2-celled sporeling with random cortical arrays and polar organiser in apical cell (D) and two 3-celled sporelings with distinct cortical arrays in apical and basal cells (E). Presented are Z-projections and central planes in XY (’). Arrows indicate the division planes. Scale bars, 10 μm.

Next, we determined if the basal position of the polar organisers was retained by the mitotic spindle and guided the plane of cytokinesis. At mitosis, roughly 30 h after plating, the polar organiser and perinuclear arrays disappeared (<5 min) and the mitotic spindle formed (5 min in Fig. 4B). The spore then passed through metaphase, anaphase, and telophase in the next 20 to 25 min. Importantly, the axis between the mitotic spindle poles was parallel to the axis formed by the polar organisers. During cytokinesis, roughly 31 h after plating, a phragmoplast formed of loosely packed, short parallel microtubules expanded centrifugally until it touched the cell cortex (0 to 40 min in Fig. 4C). The orientation of expansion was perpendicular to the mitotic spindle axis, dividing the cell asymmetrically. As the phragmoplast dismantled, cortical microtubules polymerised at distinct points across the cortex until the surface was covered by a random network (80 min in Fig. 4C). Overall, the location of the docked polar organiser defines the basal pole, determines the site of mitosis and the plane of cell division.

To investigate the microtubule organisation in the daughter cells derived from division of the spore, multi-celled sporelings were imaged at 50 h after plating. Two-celled sporelings comprised of a large apical cell and small basal cell in which microtubules were randomly arranged (Fig. 4D). Polar organisers were occasionally observed near the centroid of the apical cell. Three-celled sporelings comprised of two apical cells – in which cortical microtubules were randomly organised – and one basal cell that had initiated a tip-growing projection (rhizoid) (Fig. 4E). Microtubules were often arranged parallel to the long axis of the projection as observed in other tip-growing plant cells. Together these data indicate that microtubules re-establish after the first division to form different arrays in the apical and basal cells, reflecting their different cell identities.

### An actin filament array forms between the nucleus and the basal cortex

We hypothesised that actin filaments would also reorganise and function in the polarisation of spores. Live actin filament dynamics were visualised in spores using the *p*Mp*WDL:GFP-LifeAct* (GFP-LifeAct) reporter (Fig. S4A-C). This contains a short peptide sequence, LifeAct, that labels plant actin filaments *in vivo* (Era et al., 2009; Riedl et al., 2008; Vidali et al., 2009). In spores expressing GFP-LifeAct, a multi-layered, dense ‘mesh’ of filaments was present in the spore cell cortex at 29 h after plating (Fig. S4D, E). These filaments were randomly organised, often formed bundles and varied in length. Short interval timelapses captured the constant polymerisation and highly dynamic nature of these filaments (Fig. S4F). Taken together, GFP-LifeAct successfully labels actin filaments in spores.

To test if actin filaments reorganise during nuclear migration in spores, we simultaneously imaged actin filaments (GFP-LifeAct), the nucleus (mScarletI-N7) and the plasma membrane (mScarletI-AtLTI6b) in spores at 29 h after plating. Actin filament organisation was then compared between spores with a nucleus located at the centroid (central nucleus) and a nucleus located near the basal pole (basal nucleus). In spores with a central nucleus, bundles of actin filaments were uniformly distributed through the cortex with no actin filament arrays localised around the nucleus (Fig. 5A). By contrast, in spores with a basal nucleus, a dense array of fine actin filaments accumulated between the nucleus and the basal pole - a ‘basal actin filament array’ (Fig. 5B). Furthermore, there was a high density of actin filaments at the cell cortex below the nucleus at the basal pole – a ‘basal actin filament patch’. In timelapses, the basal actin filament array and patch were present when the nucleus was near the cell cortex and during early mitosis (Fig. 5C). In summary, a basal actin filament array and patch form when the nucleus nears the basal pole.

**Fig. 5:**
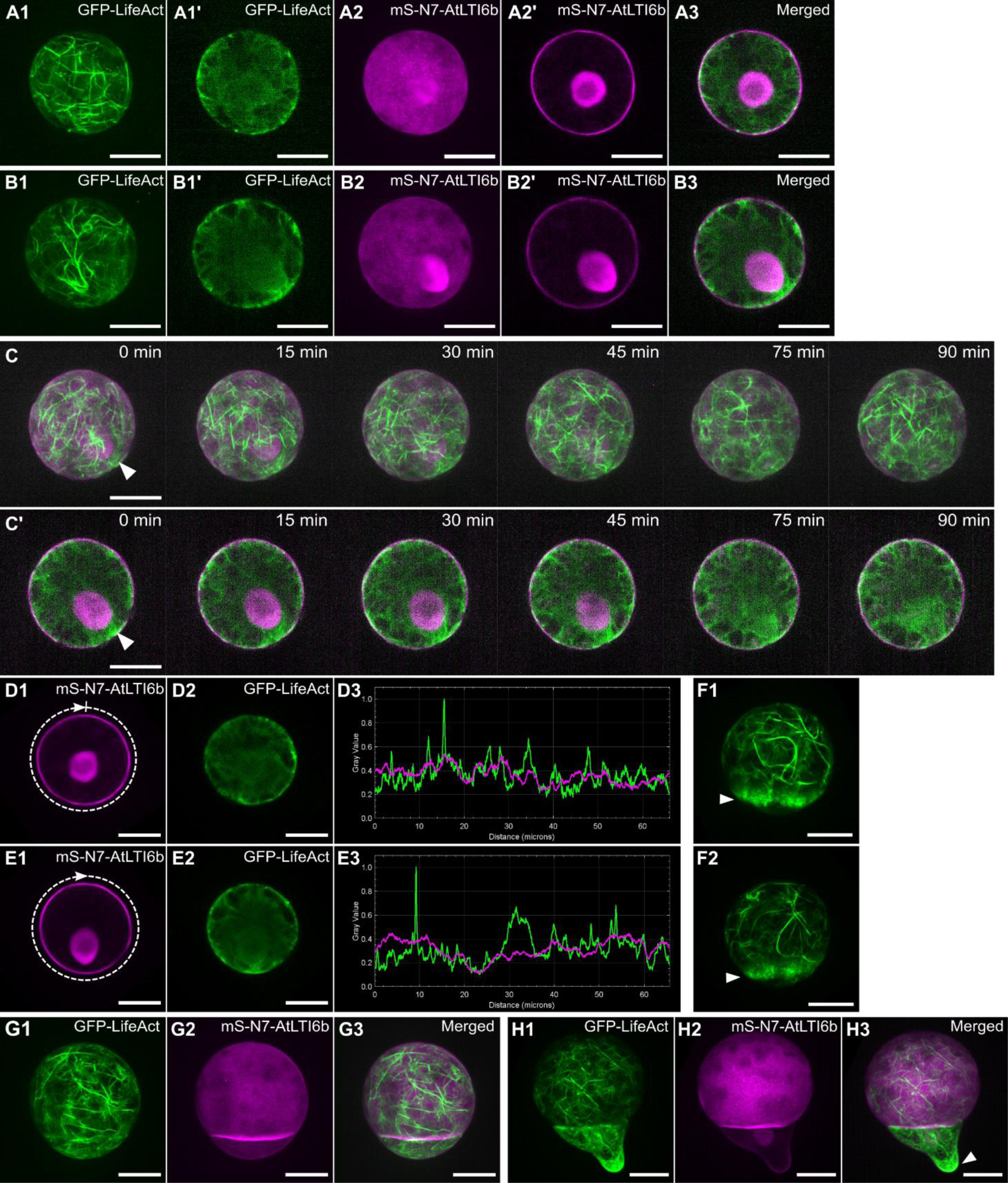
An actin filament network forms between the nucleus and basal cortex. (A, B) Organisation of actin filaments (green) in spores with a nucleus (magenta) either at the cell centre (A) or basal pole (B). Spores are expressing *p*Mp*WDL:GFP-LifeAct* and *p*Mp*UBE2:mScarletI-*At*LTI6b - p*Mp*ROP:mScarletI-N7* at 29 h after plating. Presented are Z-projections of GFP (1) and mScarlet (2) and central XY slices of GFP (1’) and mScarlet (2’). Scale bars, 10 μm. (C) Timelapse of actin filament organisation in a spore expressing *p*Mp*WDL:GFP-LifeAct* (green) and *p*Mp*UBE2:mScarletI-*At*LTI6b - p*Mp*ROP:mScarletI-N7* (magenta) starting at 29 h after plating. Presented are merged channels as Z-projections or central slices in XY (’). A white arrow indicates the actin network between the nucleus and basal cortex. Scale bars, 10 μm. (D, E) Comparison of the RFP (plasma membrane labelled by *p*Mp*UBE2:mScarletI-*At*LTI6b*) and GFP (actin filaments labelled by *p*Mp*WDL:GFP-LifeAct*) signal intensities in spores with a central nucleus (D) and basal nucleus (E) at 29 h after plating. Presented are sum-of-slices projections of the central 2.6 μm section in RFP (1) and GFP (2) used for analysis. Plots (3) present the normalised intensity values of GFP (green) and RFP (magenta) around the spore perimeter, μm. Arrows in 1 indicate the start point and direction that intensity was measured. Scale bars, 10 μm. (F) Actin filament organisation during cytokinesis of spores expressing *p*Mp*WDL:GFP-LifeAct* at 29 h after plating. Presented are Z-projections of two spores. White arrows indicate the phragmoplast. Scale bars, 10 μm. (G, H) Actin filament organisation in two-celled spores expressing *p*Mp*WDL:GFP-LifeAct* (green) and *p*Mp*UBE2:mScarletI-*At*LTI6b - p*Mp*ROP:mScarletI-N7* (magenta) captured at 29 h (G) or 50 h (H) after plating. Presented are Z-projections of the GFP (1), RFP (2) and merged channels (3). A white arrow indicates the actin network at the rhizoid cell tip. Scale bars, 10 μm.

To verify the presence and evaluate the variability of the basal actin filament patch, the GFP (GFP-LifeAct) and RFP (mScarletI-AtLTI6b) fluorescence intensities around the cell surface were quantified and compared. In spores with a central nucleus, GFP and RFP intensities varied together and their intensity ratio was stable around the entire cell circumference (Fig. 5D). In spores with a basal nucleus, the GFP intensity was consistently greater than the RFP intensity over an approximately 10 µm region located half-way around the spore circumference - between 28 and 38 µm distance - which included the basal pole (Fig. 5E). This increased GFP to RFP ratio in the basal cortex was consistent among all spores quantified with a basal nucleus (Fig. S5). This verified our observations that actin filaments only accumulated at the basal pole when the nucleus was basally positioned. However, the intensity and appearance of the GFP peak was variable between spores, appearing as one wide peak, as two high separate narrower peaks, or as multiple smaller scattered peaks at the basal pole (Fig. S5). This suggests that the basal actin filament network is highly dynamic. In summary, a dense network of fine actin filaments forms at the basal cortex as the nucleus migrates from the cell centroid to basal pole.

### A dense actin filament network develops in the basal daughter cell after cell division

Next, we determined if actin filaments reorganise during or after the first asymmetric cell division. In spores undergoing cytokinesis at 29 h after plating, a short, dense array of actin filaments corresponding to the phragmoplast developed (Fig. 5F). On the apical side of the phragmoplast, actin filaments were organised into bundled arrays but not on the basal side. After cytokinesis, actin filaments were bundled in the apical cell and relatively unbundled fine arrays in the basal cell (Fig. 5G). In two-celled spores at 50 h after plating, the basal cell had elongated at the apex and an array of fine actin filaments developed at the site of tip-growth (Fig. 5H). In summary, actin filament organisation differs between the apical and basal hemisphere of the spore during and after cytokinesis.

### Depolymerisation of actin filaments decreases division asymmetry

To test the hypothesis that actin filaments are required for the basal migration of the nucleus leading to the first asymmetric cell division, actin filaments were depolymerised in sporelings using Latrunculin B (LatB) and the symmetry of the cell division was measured. To identify a LatB dose at which actin filaments would be depolymerised but cell division could still proceed, wild type spores were grown on media containing a range of LatB concentrations for 4 days (Fig. S7). On 0.1% DMSO (control), most spores divided to produce a rhizoid cell and proliferating apical cell. On 0.03 µM LatB, spores divided but lacked rhizoids. On 0.1 µM LatB, some spores divided to form two equal sized chloroplast-filled cells. These data indicate that LatB treatment blocked rhizoid outgrowth and disrupted cell division.

To test if LatB disrupted the asymmetry of the first cell division, the position of the new cell wall was defined in LatB-treated sporelings. Developing sporelings expressing reporters labelling actin filaments, the plasma membrane and the nucleus were grown on 0.1% DMSO (control) or 0.1 µM LatB for 48 h. Actin filaments were present at the cortex of DMSO-treated sporelings but not of LatB-treated sporelings, confirming that actin filaments were depolymerised by the 0.1 µM LatB treatment (Fig. 6A-C). On the DMSO control, 96% of sporelingss – in which the division plane could be determined – divided asymmetrically and only 4% divided symmetrically (Fig. 6A, D). By comparison, on 0.1 µM LatB, 78% of spores divided asymmetrically and 22% divided symmetrically (Fig. 6B-D). The increase in the percentage of symmetric divisions on LatB-treatment compared to control treatments indicates that depolymerisation of actin filaments disrupts cell division asymmetry. In the absence of actin filaments, the nucleus remains in the cell centre leading to a symmetric cell division in some cells. These data are consistent with the hypothesis that actin filaments are required for the basal migration of the nucleus to define the asymmetry of the first division.

**Fig. 6:**
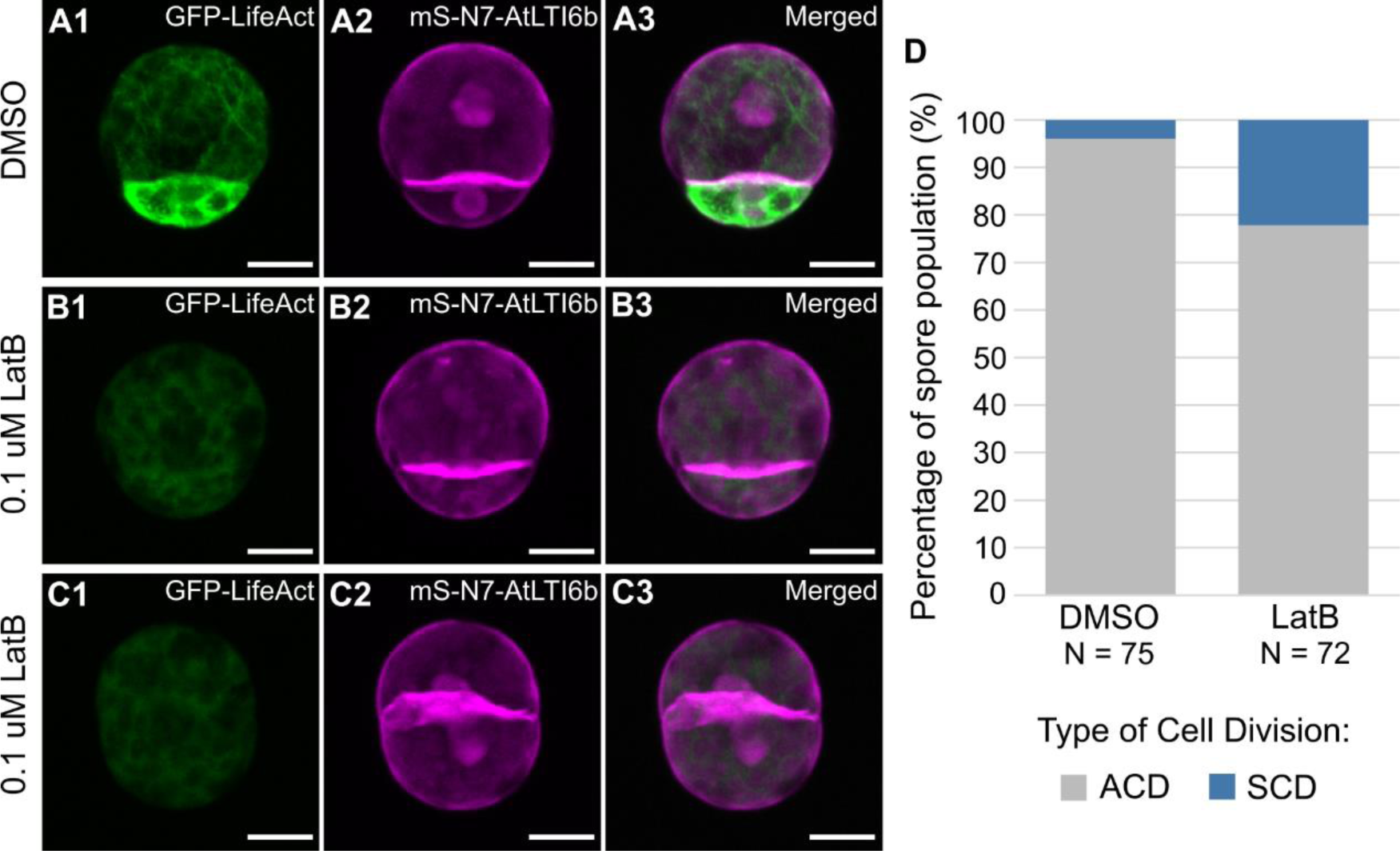
Depolymerisation of actin filaments increases symmetric cell divisions. (A -C) Plasma membrane (magenta) and actin filaments (green) showing the division plane of spores grown on 0.1% DMSO (A) or 0.1 μM LatB (B, C) for 48 h. Spores express *p*Mp*WDL:GFP-LifeAct* and *p*Mp*UBE2:mScarletI-*At*LTI6b - p*Mp*ROP:mScarletI-N7*. Presented are Z-projections of GFP (1), mScarlet (2) and merged (3). Scale bars, 10 μm. (D) Percentage of spores that divided asymmetrically (ACD) and symmetrically (SCD) after 48 h growth on 0.1% DMSO and 0.1 μM LatB. N is the number of spores scored.

## DISCUSSION

Cellular asymmetry develops in the *M. polymorpha* spore and directs nuclear movement to orient the first asymmetric cell division plane. At 24 h after plating, the nucleus is located at the spore centroid but migrates to the cortex at the basal pole over the course of less than an hour. A dense network of fine actin filaments is located between the nucleus and the basal cortex. As actin-myosin systems often provide a mechanical force for organelle movement, we speculate that this actin filament network pulls the nucleus to the basal pole. Subsequently the mitotic spindle forms near the basal pole. The phragmoplast then expands at the mitotic spindle equator, dividing the spore into a relatively large apical cell and small basal cell. Therefore, nuclear migration establishes an internal apical-basal asymmetry that directs the first division of the spore.

We hypothesised that filamentous actin was required for nuclear movement and to define the asymmetry of the first division. Upon depolymerisation of actin filaments, we observed a decrease in cell division asymmetry; while basal cell was approximately 10% the volume of the apical cell in control treatments (DMSO), both cells were more similar in size on LatB treatment. These data are consistent with the hypothesis that actin-mediated nucleus movement is required for the development of cellular asymmetry to orient the first asymmetric cell division in spores.

During the first 24 h of spore development, a dynamic cortical microtubule array forms in a random organisation, as theoretically expected for a spherical cell where there is no preferential axis of growth. Between roughly 28 and 32 h of spore development, and approximately 1 h before nuclear migration, a pair of microtubule organising centres - polar organisers – form on opposite sides of the nucleus. The axis between these two polar organisers rotates before and during nuclear migration. One polar organiser leads this migration by nucleating a dense astral microtubule array that polymerises towards the cortex at the basal pole. We speculate that this astral array anchors to a polar cortical domain to guide the nucleus to this domain. Simultaneously, cortical microtubules disappear from the apical hemisphere but remain near the basal hemisphere. As this basal cortical array forms in prophase, and roughly aligns with the equator of the future mitotic spindle, we speculate that these microtubules deposit proteins in a cortical division zone to guide cytokinesis.

The space between the nucleus and the basal cortex is filled with an astral microtubule array nucleated from the basal polar organiser and a network of fine filamentous actin. We speculate a model for their interaction (Fig. S8). First an initial polarity cue orients the formation of a polarity domain at the basal pole. This domain anchors the plus ends of astral microtubules initiated from the closest polar organiser. As progressively more microtubules are initiated from this basal polar organiser, a dense basal astral array forms which orients the nucleus and polar organisers towards the basal pole. Simultaneously, actin filament nucleation begins at the basal pole. The filamentous actin network polymerises inwards to attach to and pull the nucleus. This nuclear migration orients the first division to position a differentiated rhizoid cell at the basal pole and a proliferating cell at the apical pole.

## MATERIALS AND METHODS

### Reporter constructs

The *p*Mp*EF1α:GFP-*Mp*TUB1* and *p*Mp*EF1α:GFP-*At*ENDBINDING1a* reporters label microtubules and microtubule plus ends respectively (Buschmann et al., 2016).

The *p*Mp*ROP:mScarletI–N7* and *p*Mp*UBE2:mScarletI-*At*LTI6b* reporters label the nucleus and plasma membrane respectively. Cloning used the promotor of the Mp*ROP* gene and components from the OpenPlant toolkit (Sauret-Güeto *et al*., 2020; Mulvey & Dolan, 2023).

The *p*Mp*WDL:GFP-LifeAct* reporter labels actin filaments and is composed of a LifeAct sequence – ATGGGTGTCGCAGATTTGATCAAGAAAT TCGAAAGCATCTCAAAGGAAGAA - fused to GFP expressed under the promotor of the Mp*WAVE-DAMPENED-LIKE* gene (Champion et al., 2021).

### Amplification of plasmids and transformation into *Agrobacterium*

Plasmids were transformed into One Shot OmniMAX 2-T1R chemically competent *E. coli* cells (ThermoFisher Scientific) by heat-shock at 42 °C for 30 seconds then ice for 2 minutes. After adding 250 μL S.O.C medium, the mixture was incubated at 37 °C for 1 hour shaking at 225 rpm. The mixture was diluted into LB medium (1:50), spread onto solid LB media plates containing 100 μg/mL spectinomycin and incubated overnight at 37 °C. Single colonies were grown at 37 °C overnight in liquid LB media containing 100 μg/mL spectinomycin. Plasmids were isolated from the transformed *E. coli* cells using the GeneJet Plasmid MiniPrep kit (ThermoFisher Scientific).

Amplified plasmids were transformed into the *Agrobacterium tumefaciens* GV3101 strain by electric shock. 1 μL plasmid to 50 μL *Agrobacterium* was micro-pulsed then 1 mL S.O.C media added. Transformed *Agrobacterium* was grown for 2 days at 28 °C on LB media plates containing 50 μg/mL gentamycin, 50 μg/mL rifampicin, and 50 μg/mL spectinomycin. single transgenic colonies transferred into liquid LB antibiotic media.

### Transformation of Marchantia polymorpha

Plasmids were transformed into wild type *M. polymorpha* sporelings – derived from crossing Tak-1 and Tak-2 - using transgenic *Agrobacterium* following the method developed in Ishizaki *et al*., 2008 and improved upon in Honkanen *et al*., 2016.

### Plant lines, growth conditions and crossings

The wild type *Marchantia polymorpha* accessions used were Takaragaike-1 (Tak-1) male and Takaragaike-2 (Tak-2) female. Plants were grown on ½-strength B5 Gamborg’s medium containing 1.5 g/L B5 Gamborg, 0.5 g/L MES hydrate, 1% sucrose, pH adjusted to 5.5, set with 1% agar. Plants were grown at 23 °C in continuous white light at 50 - 60 μmol m²s¹.

To induce reproductive development, plants were potted on soil, containing a 1:3 ratio of fine vermiculite and Neuhaus N3 compost, within SacO2 Microbox containers and grown 20°C in long day conditions of 16 h light, 8 h dark. White light was set at 50 - 60 μmol m²s¹ and enhanced with far-red light at 30 - 40 μmol m²s¹. Male and female plants were crossed to generate spores. Sporangium were sterilised in 1 % sodium dichloroisocyanurate (NaDCC) for 3 min before washing and bursting in sterile water.

### Drug treatments

Wild type spores were grown on Gamborg media plates containing Oryzalin or Latrunculin B dissolved in DMSO. The final oryzalin concentrations were 0.01 µM, 0.03 µM, 0.1 µM, 0.33 µM, 1 µM, 3.3 µM and 10 µM. The final Latrunculin B concentrations were 0.01 µM, 0.03 µM, 0.1 µM, 0.33 µM, 1 µM and 5 µM. 0.1% DMSO was used as a control.

Spores from a cross between plants expressing *p*Mp*WDL:GFP-LifeAct* and *p*Mp*ROP:mScarletI-N7 - p*Mp*UBE2:mScarletI-*At*LTI6b* were grown on a Gamborg media slab containing 0.1% DMSO or 0.1 µM Latrunculin B within imaging chambers.

### Stereomicroscope imaging

Spores grown on media plates were imaged with a Keyence VHX-7000 digital microscope equipped with a VHX-7020 camera and VH-ZST lens.

### Spinning disk imaging

Imaging chambers were made following the protocol in Kirchhelle & Moore, 2017 with a few adjustments. A breathable gum boarder (Carolina Observation gel) was filled with Gamborg media and layered with cellophane soaked in liquid Gamborg media (½-strength B5 Gamborg’s medium without agar). One sterile sporangium was burst into 200 µL water and 40 µL of the solution added to the chamber. Spores were grown within the chamber for 29 to 50 h before imaging.

Imaging used an Olympus IX3 Series (IX83) inverted microscope equipped with a Yokogawa W1 spinning disk, Hamamatsu ORCA-Fusion CMOS camera and a 100x/1.45 NA oil objective or 40x/0.75 NA air objective. For GFP, excitation was set at 488 nm and emission captured at 525 nm. For mScarlet, excitation was set at 561 nm and emission captured at 617 nm. For chlorophyll, excitation was set at 640 nm and emission captured at 685 nm. Z-stacks of 20 to 22 µm with 0.26 µm slices. Lazer power and exposure time were tested and adjusted to prevent bleaching overtime. A Z-Drift compensation autofocus system stabilised imaging overtime. As spore populations developed asynchronously, and were heterogeneous for the reporter expression, specific spores within each population were selected for imaging.

A few alterations to this method were required to capture microtubule plus ends (GFP-AtEB1). A Piezo Z stage and Hamamatsu ORCA-Flash 4.0 camera were used to rapidly capture Z-stacks of 0.99 µm with 0.33 µm slices at 1.2 s intervals.

Gemmae expressing GFP-LifeAct were setup and grown in imaging chambers for 1 day and imaged using an Olympus IX3 Series (IX83) inverted microscope equipped with a Yokogawa W1 spinning disk, Hamamatsu ORCA-Fusion CMOS camera and 10x/0.4 NA air and 100x/1.45 NA oil objectives. Excitation set at 488 nm and emission captured at 525 nm.

### Image deconvolution and image analysis

Image deconvolution used Huygens software (Scientific Volume Imaging). Z-projections and central slices were converted using ImageJ Fiji (Schindelin et al., 2012). For temporal projections, each image slice was deconvolved in Huygens then maximum projections for each timepoint were generated in ImageJ Fiji before temporal projection in ImageJ Fiji.

Analysis of cortical microtubule density used the ImageJ LPX package published in Higaki *et al*., 2010. From Z-projections, the spore perimeter was outlined in ImageJ Fiji at each timepoint. Microtubules were skeletonised using the LPX Filter2d with the Otsu method and a line extract value of 5. This image was masked by the spore outline, and the skeletonised microtubules were analysed using the LPX script. Graphs were created in Microsoft Excel.

The signal intensity profiles of GFP and mScarlet fluorophores around the spore perimeter were generated in ImageJ Fiji, using a script written by Tomas Lendl from the BioOptics Facility at the Vienna BioCenter. A sum-of-slices projection of a 5.2 µm region surrounding the medial plane was created for each spore. The perimeters were outlined using the segmented line tool set at spline 5, starting at 12 o’clock and moving clockwise. The script then applied background subtraction, scaled the two fluorophore intensities to the mean, and normalised the relative signal intensities. The resultant fluorophore intensity profiles, ranging from 0 to 1, were plotted against the distance around cell perimeter (µm) from the starting point. Plots were automatically generated using the ImageJ Fiji script.

### Drug treatments on spores

Spores were grown on Gamborg media plates containing oryzalin or Latrunculin B dissolved in DMSO. The final oryzalin concentrations were 0.01 µM, 0.03 µM, 0.1 µM, 0.33 µM, 1 µM, 3.3 µM and 10 µM. The final Latrunculin B concentrations were 0.01 µM, 0.03 µM, 0.1 µM, 0.33 µM, 1 µM and 5 µM. 1% DMSO was used as a control.

## ACKNOWLEDGEMENTS

We thank Katharina Jandrasits and Magdalena Mosiolek for their lab support. We acknowledge the BioOptics Facility and Plant Science Unit at the Vienna BioCenter Core Facilities, Austria. In particular, we thank Pawel Pasierbek and Alberto Monerono Cencerrado for their imaging expertise and Thomas Lendl for producing the image analysis script. Maria Gravato-Nobre and Clement Champions for their mentoring and great discussions. Charlotte Kirchhelle for sharing the imaging chamber design and discussing experiments. We are grateful to Henrik Buschmann for insightful discussions and sharing reporter lines.

## DATA AVALIABILITY

All data are available.

## SUPPLEMENTARY FIGURES

**Fig. S1:**
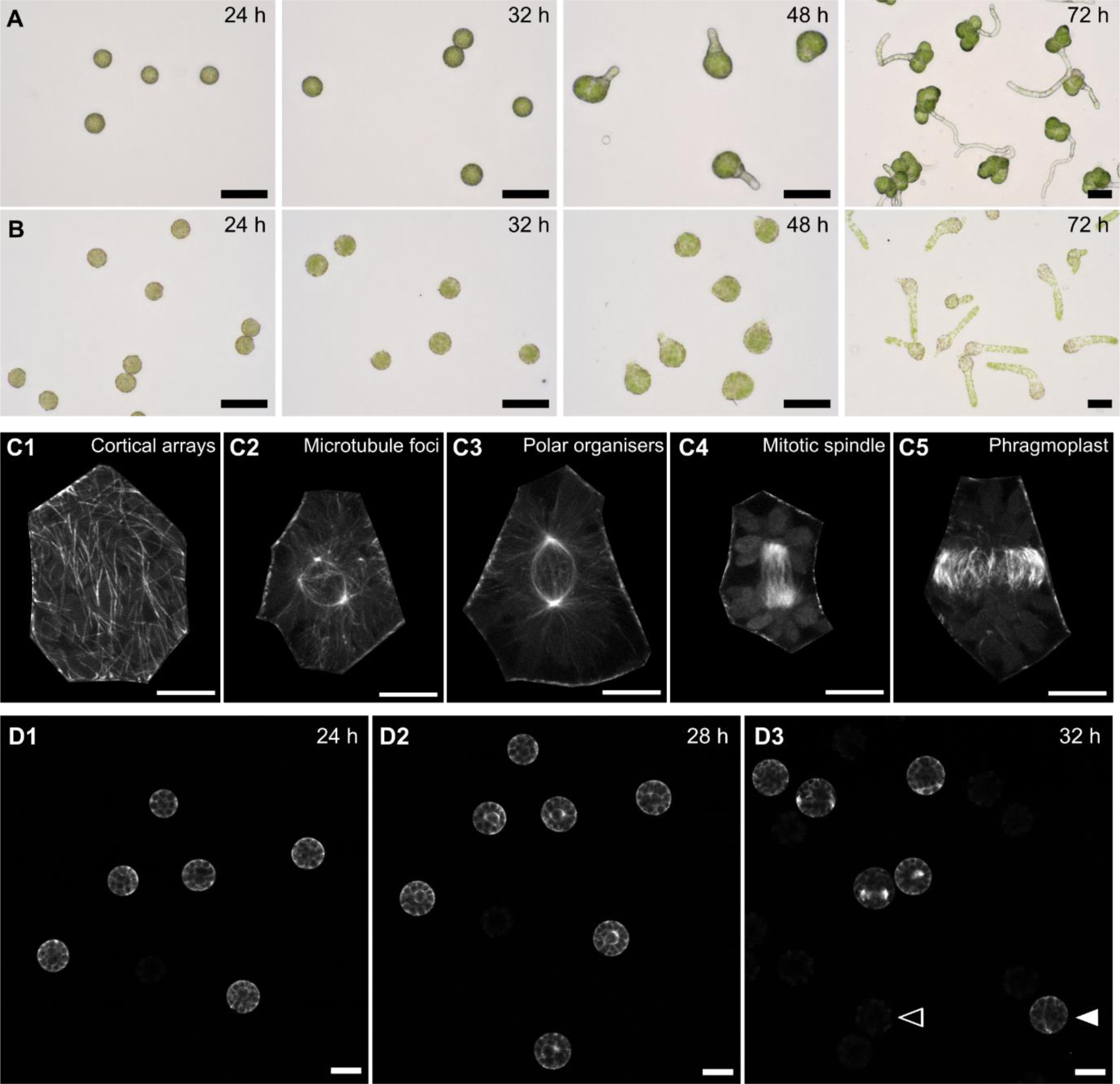
Development of wild type spore populations and the microtubule structures present in *M. polymorpha* cells. (A, B) Development of wild type spore populations grown on media plates (A) and within an imaging chamber (B). Imaged at 24, 32, 48 and 72 h after plating. Presented are different spores from the same population. Scale bars, 50 μm. (C) The microtubule organisations in wild type epidermal cells of 2-day-old gemmae expressing *p*Mp*EF1α:GFP-*Mp*TUB1.* Presented are Z-projections of individual cells with surrounding cells removed. Scale bars, 10 μm. (D) Population of spores expressing *p*Mp*EF1α:GFP-*Mp*TUB1* imaged at 24, 28 and 32 h after plating. The population is mixed; half express the reporter (filled arrow) and half do not (empty arrow). Presented are central slices in XY. Scale bars, 20 μm.

**Fig. S2:**
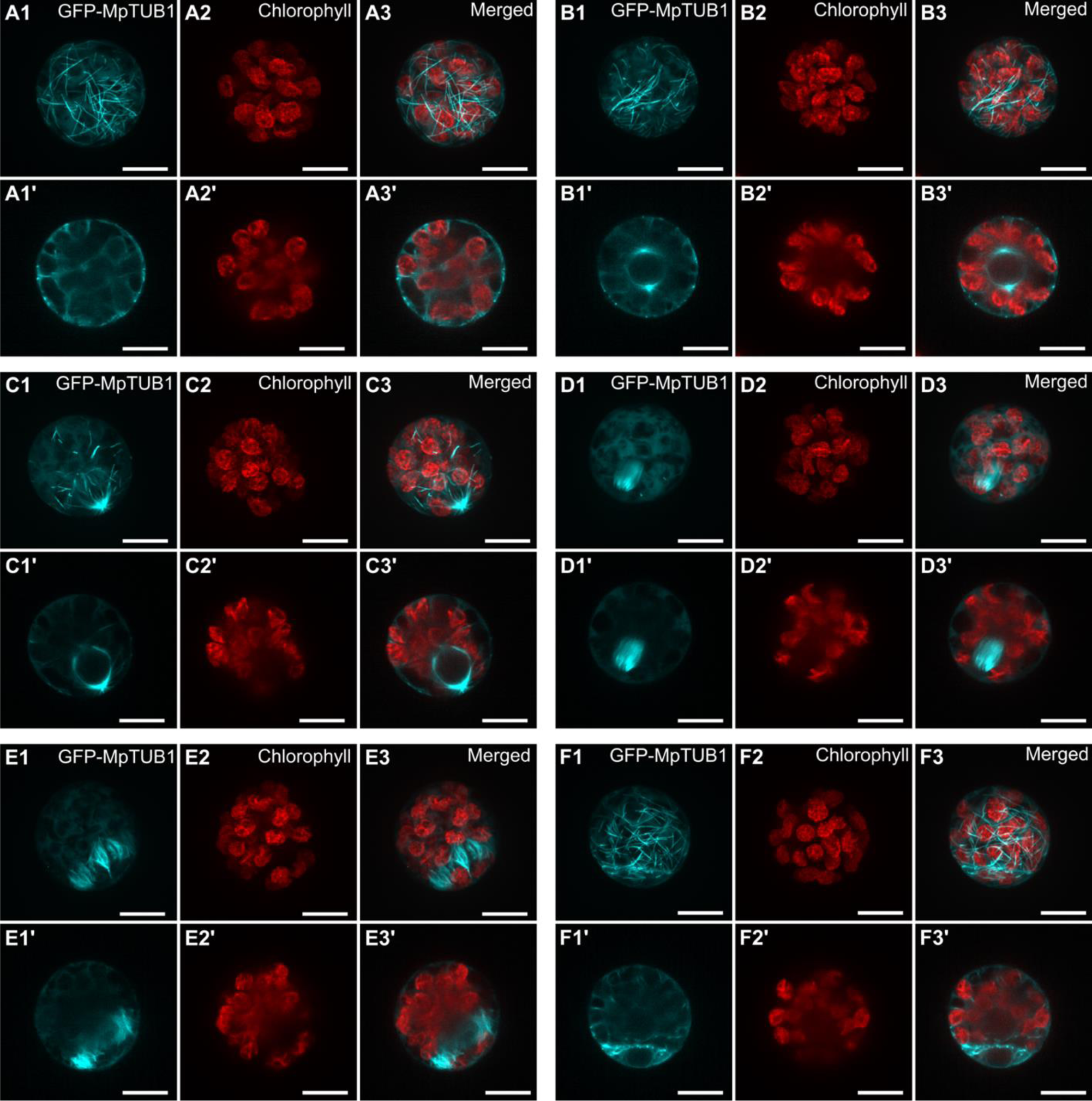
Chloroplasts distribute asymmetrically in polarised spores. Organisation of microtubules and chloroplasts in spores expressing *p*Mp*EF1α:GFP-*Mp*TUB1* at 29 h to 32 h after plating. Shown are GFP-MpTUB1 fluorescence (cyan, A1-F1), chlorophyll autofluorescence (red, A2-F2) and merged images (A3-F3) of each spore. Presented are the Z-projections (A-F) and central planes in XY (A’-F’) of each channel. (A) Spores with no polar organisers are packed with chloroplasts. (B) Spores with two central polar organisers have a central clear zone. (C) Spores with a basal microtubule basket and (D) spores with a late mitotic spindle/early phragmoplast have a basal zone clear of chloroplasts. (E) Spores undergoing cytokinesis and (F) divided spores have an apical zone/cell with many chloroplasts and basal zone/cell with few chloroplasts. Scale bars, 10 μm.

**Fig. S3:**
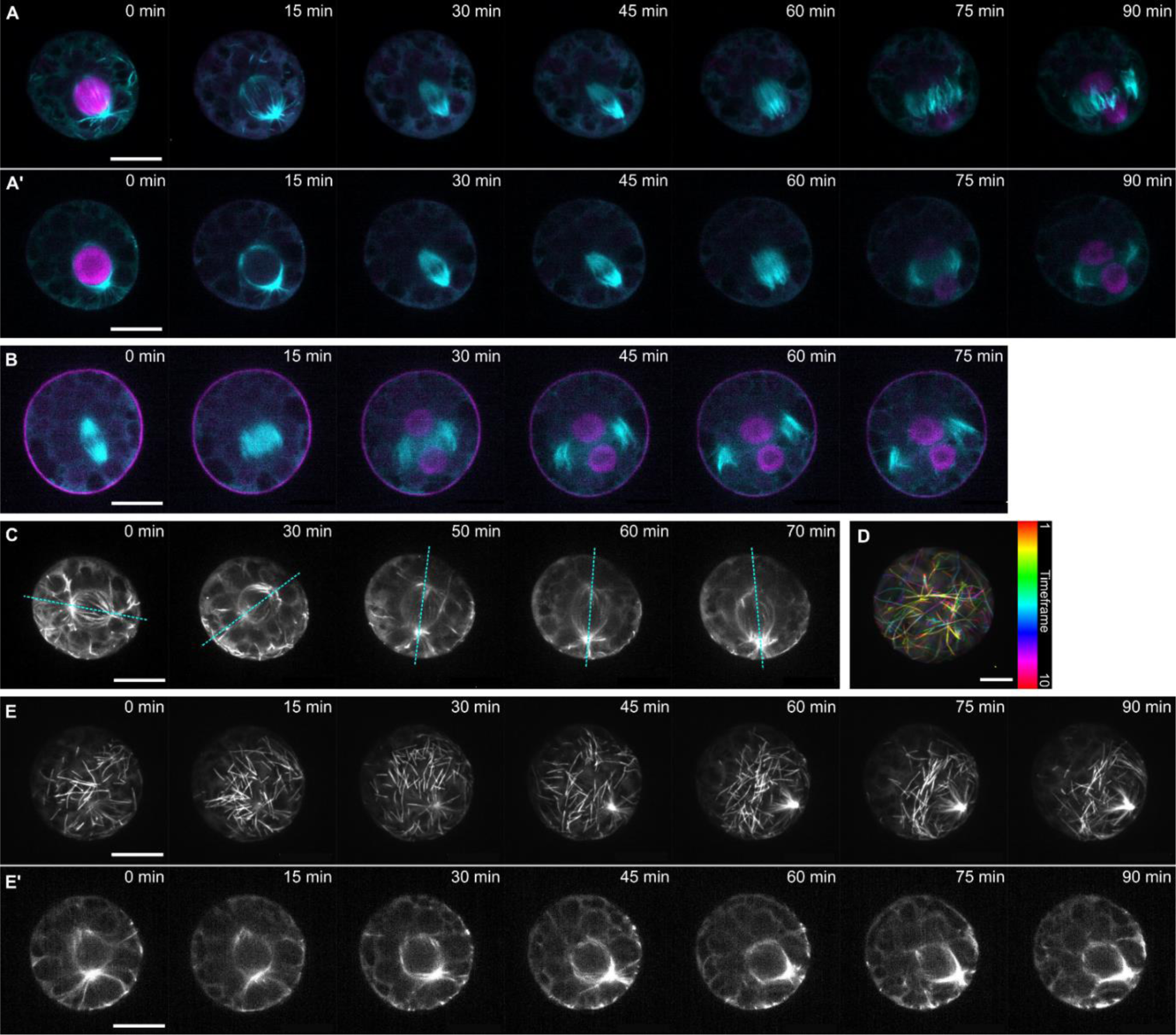
The nuclear localised signal disappears on mitosis and reappears in the two daughter cells. (A, B) Timelapses of dividing spores expressing *p*Mp*EF1α:GFP-*Mp*TUB1 (*cyan) and *p*Mp*ROP:mScarletI-N7* - *p*Mp*UBE2:mScarletI-*At*LTI6b* (magenta) at 29 h after plating. Presented are Z-projections (A) and central planes in XY (A’, B). Scale bars, 10 μm. (C) Timelapse of spore expressing *p*Mp*EF1α:GFP-*Mp*TUB1* at 29 h after plating. Presented are Z-projections of central slices showing rotation of polar organisers in 3 dimensions. Cyan lines indicate the polar organiser axis. Scale bar, 10 μm. (D) Temporal projection of cortical microtubules in a spore expressing *p*Mp*EF1α:GFP-*Mp*TUB1* at 29 h after plating. Time intervals of 20 s. Scale bar, 10 μm. (E) Timelapse of a spore expressing *p*Mp*EF1α:GFP-*Mp*TUB1* at 29 h after plating. Presented are Z-projections showing the depletion of cortical microtubules from the apical cortex as the polar organiser, seen in the central XY planes (’), migrates to the basal pole. Scale bars, 10 μm.

**Fig. S4:**
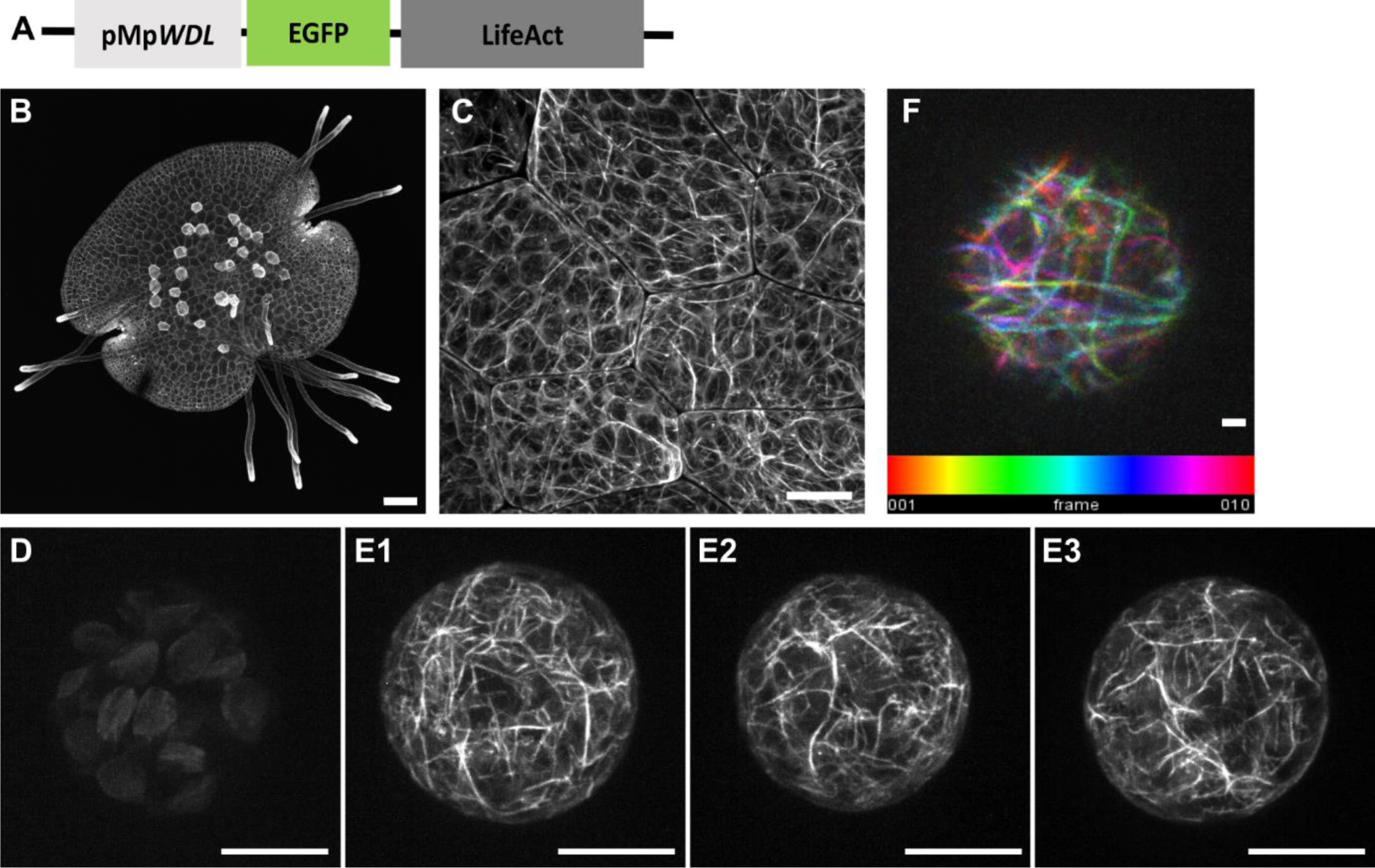
GFP-LifeAct labels dynamic cortical actin filaments in *M. polymorpha* spores. (A) Schematic of the actin filament reporter, *p*Mp*WDL:GFP-LifeAct*. (B) Actin filaments are labelled in all the cells of a 1-day-old gemma expressing *p*Mp*WDL:GFP-LifeAct*, strongly at the rhizoid tips. Scale bar, 100 μm. (C) Actin filaments are labelled in the cortex of epidermal cells from a 1-day-old gemma expressing *p*Mp*WDL:GFP-LifeAct*. Scale bar, 10 μm. (D) GFP autofluorescence of a wild type spore at 29 h after plating. Scale bars, 10 μm. (E) Actin filaments are labelled in the cortex of spores expressing *p*Mp*WDL:GFP-LifeAct* at 29 h after plating. Scale bars, 10 μm. (F) Temporal projection of actin filaments polymerising across the cortex of a spore expressing *p*Mp*WDL:GFP-LifeAct* at 30 h after plating. Timelapse captured the top 5 μm of the spore with 3 s intervals. Z-projections were generated before temporal projection. Scale bar, 1 μm.

**Fig. S5:**
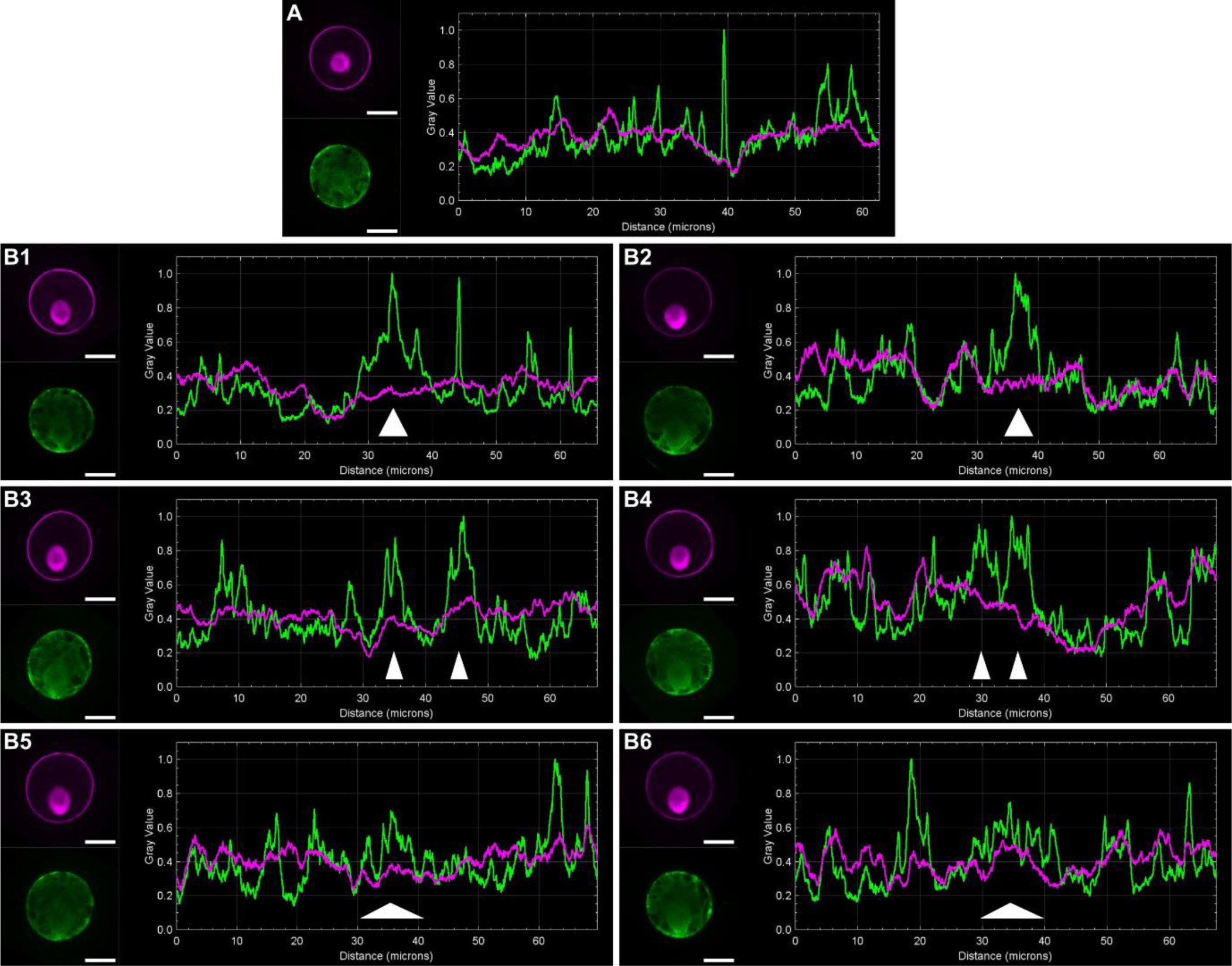
Variability in the intensity and area of the actin filament network at the basal pole. (A, B) Further comparison of the RFP (plasma membrane labelled by *p*Mp*UBE2:mScarletI-* At*LTI6b*) and GFP (actin filaments labelled by *p*Mp*WDL:GFP-LifeAct*) signal intensities in a spore with a central nucleus (A) and six spores with a basal nucleus (B) at 29 h after plating. Images present sum-of-slices projections of the central 2.6 μm section in RFP (magenta) and GFP (green) used for analysis. Plots present the normalised intensity values of GFP and RFP around the spore perimeter, μm, proceeding clockwise from 12 o’clock. Regions of increased GFP signal at the basal pole are indicated by white arrows. Scale bars, 10 μm.

**Fig. S6:**
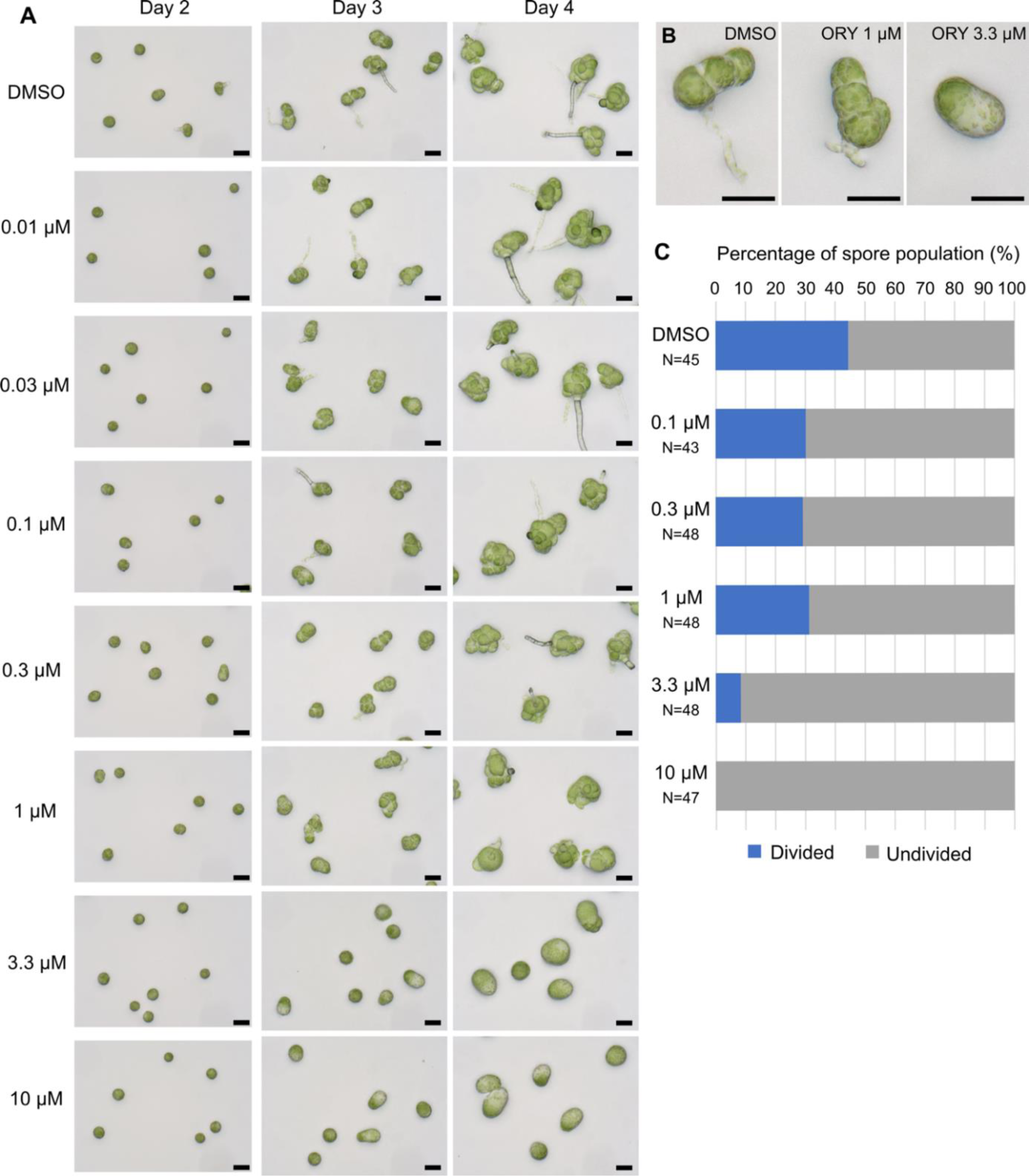
Oryzalin inhibits spores division and promotes swelling. (A) Development of a wild type spore population at 2, 3 and 4 days after plating on increasing doses of oryzalin or a DMSO control. Scale bars, 50 μm. (B) Enlarged images of representative spores grown on DMSO, 1 μM and 3.3 μM oryzalin for 3 days. Scale bars, 50 μm. (C) Percentage of divided (blue) and undivided (grey) spores in a population grown for 2 days on 0.1 μM, 3.3 μM and 10 μM oryzalin and DMSO. N is the number of spores quantified for each dose.

**Fig. S7:**
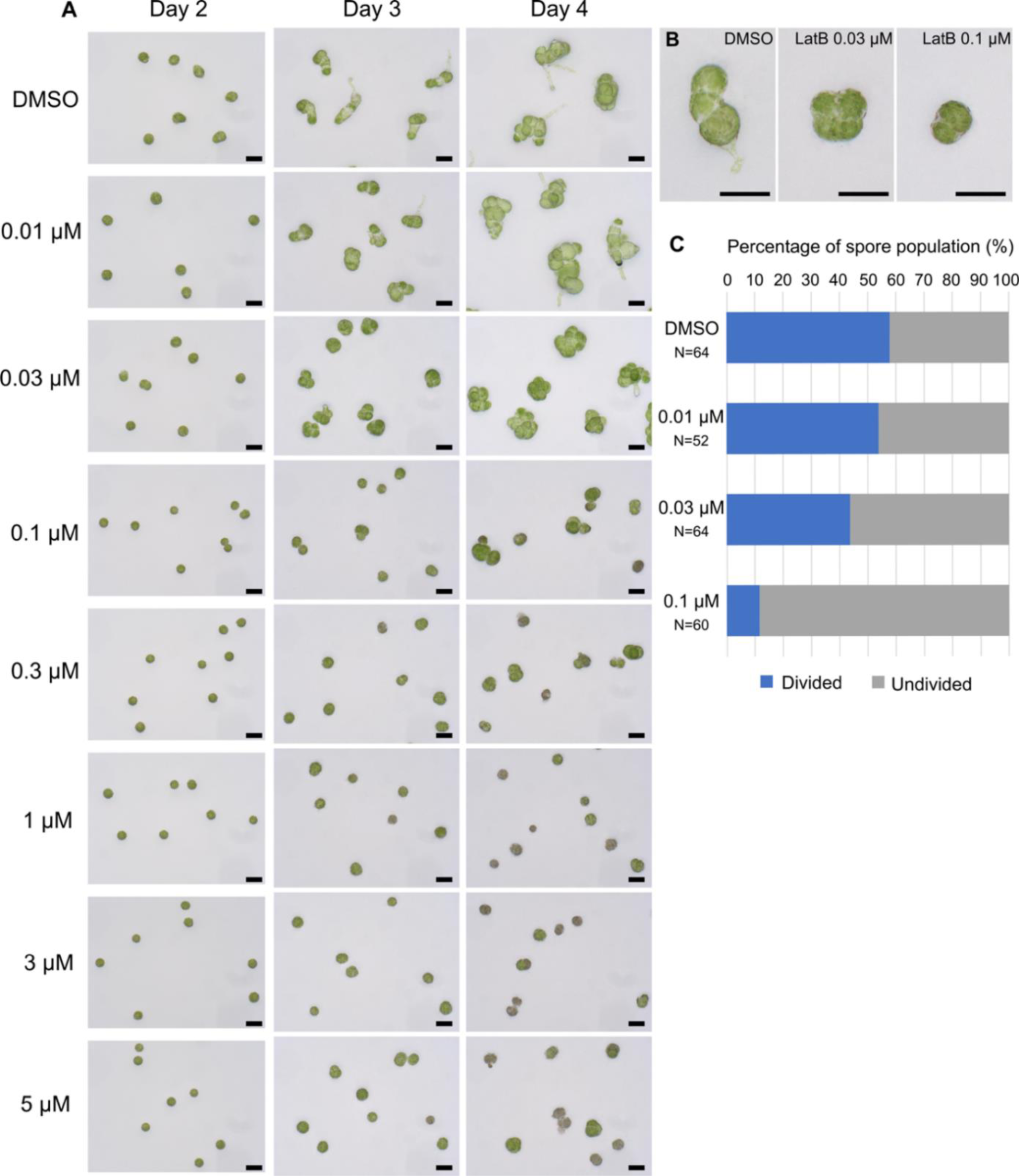
Latrunculin B disrupts rhizoid formation and appears to induce more symmetrical divisions in spores. (A) Development of a wild type spore population at 2, 3 and 4 days after plating on increasing doses of LatB or a DMSO control. Scale bars, 50 μm. (B) Enlarged images of representative spores grown on DMSO, 0.03 μM and 0.1 μM LatB for 3 days. Scale bars, 50 μm. (C) Percentage of divided and undivided spores in populations grown on 0.01 μM, 0.03 μM and 0.1 μM LatB and DMSO for 2 days. N is the number of spores quantified.

**Fig. S8:**
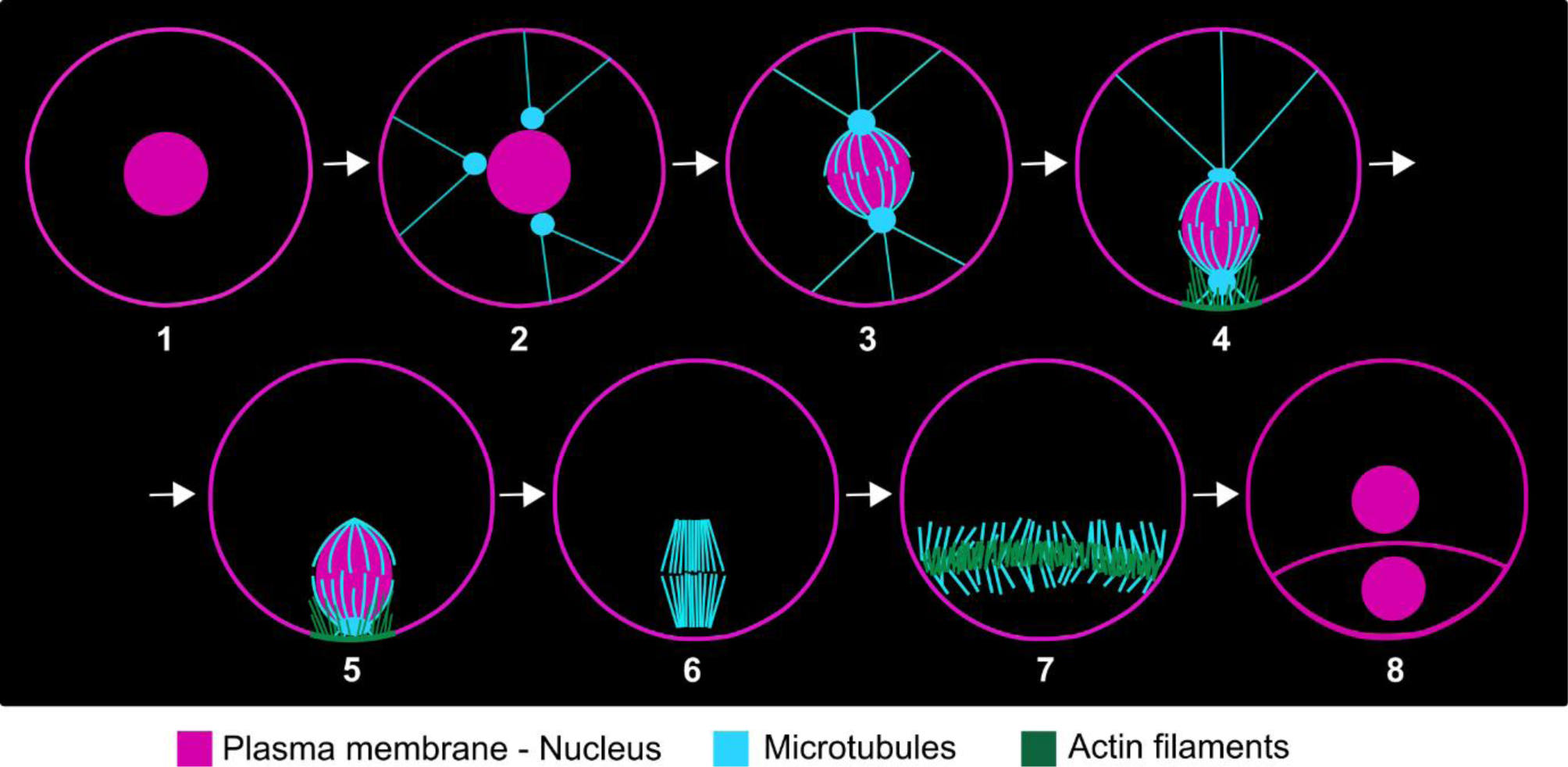
Current model of cytoskeleton reorganisation and nuclear migration during spore polarisation and the first asymmetric cell division.

**Movie 1: Tracking of microtubule polymerisation across the cortex using EB1** Movie of EB1 comets moving across the cortex of a spore expressing *p*Mp*EF1α:GFP-* At*EB1a* at 29 h after plating. Presented are deconvolved Z-projections of slices at the cell surface, taken at 1.2 s time intervals.

**Movie 2: Tracking of microtubule polymerisation from the basal polar organiser using EB1** Movie of EB1 comets moving away from the basal pole organiser (positioned at the top of the cell) in a spore expressing *p*Mp*EF1α:GFP-*At*EB1a* at 29 h after plating. Presented are deconvolved Z-projections of slices at the cell surface, taken at 1.2 s time intervals.

